# Increased rate of *de novo* single nucleotide mutation in house mice born through assisted reproduction

**DOI:** 10.1101/2025.06.27.662069

**Authors:** Laura Blanco-Berdugo, Alexis Garretson, Beth L. Dumont

**Author notes:** **Address for Correspondence**: Beth Dumont, The Jackson Laboratory 600 Main Street, Bar Harbor, ME 04609, E. **Author Contributions:** This project was conceptualized and designed by BLD and LBB. LBB performed all experimental investigation and led formal analysis, with supervision and funding support from BLD. BLD and AG contributed to formal analysis and data visualization. LBB and BLD wrote the original manuscript draft, with substantial input and review from AG. **Competing Interest Statement:** The authors have no competing interests to disclose. **Classification**: Biological Sciences; Genetics.

## Abstract

Approximately 2.6% of live births in the United States are conceived using assisted reproductive technologies (ARTs). While some ART procedures, including in vitro fertilization (IVF) and intracytoplasmic sperm injection, are known to alter the epigenetic landscape of early embryonic development, their impact on DNA sequence stability is unclear. Here, we leverage the strengths of the laboratory mouse model system to investigate whether a standard ART regimen—ovarian hyperstimulation, gamete isolation, IVF, embryo culture, and embryo transfer—affects genome stability. Age-matched cohorts of ART-derived and naturally conceived C57BL/6J inbred mice were reared in a controlled setting and whole genome sequenced to ∼50x coverage. Using a rigorous pipeline for *de novo* single nucleotide variant (dnSNV) discovery, we observe a ∼30% increase in the dnSNV rate in ART-compared to naturally-conceived mice. Analysis of the dnSNV mutation spectrum identified signature contributions related to germline DNA repair activity, affirming expectations and evidencing the quality of our dnSNV calls. We observed no enrichment of dnSNVs in specific genomic contexts, suggesting that the observed rate increase in ART-derived mice is a general genome-wide phenomenon. Similarly, we show that the developmental timing of dnSNVs is similar in ART- and natural-born cohorts. Together, our findings show that ART is moderately mutagenic in house mice and motivate future work to define the precise procedure(s) associated with this increased mutational vulnerability. While we caution that our findings cannot be immediately translated to humans, they nonetheless emphasize a pressing need for investigations on the potential mutagenicity of ART in our species.

**SIGNIFICANCE STATEMENT:** This study investigates whether assisted reproductive technologies (ARTs) increase the risk of inherited genetic mutations in offspring. Using a well-controlled mouse model system, we compared the de novo mutation burden in genomes of mice conceived through ART to a naturally conceived cohort. We find a ∼30% increase in new DNA mutations in ART-conceived mice, suggesting that ART procedures have a genome destabilizing effect. This increase in mutation rate appears to be uniform across the genome, rather than attributable to specific genomic contexts. While we caution against the direct translation of our findings to humans, our work nonetheless highlights the need for further research into the genetic safety of ART in people.

## INTRODUCTION

Worldwide, approximately 1,000,000 babies are born by assisted reproductive technologies (ART) each year (1), accounting for over 2.5% of live-births in the United States (2) and many other Western countries (3, 4). This number has been steadily increasing over the last decade (5) and is expected to continue to increase in the face of rising infertility rates (6) and increases in the age at reproduction (7). The human health consequences of an increased reliance on ARTs are potentially substantial. Prior work has shown that various ART procedures are associated with altered methylation patterns in both model organisms (8–11) and humans (12–17), with downstream consequences for the expression of key genes required for totipotency and early development (18–22). These molecular changes are associated with increased rates of imprinting related disorders (23, 24), and may contribute to higher rates of adverse pregnancy, perinatal, neonatal, and long-term health outcomes in ART-derived offspring (25–28).

While it is well-established that various ARTs can perturb the dynamics of epigenetic reprogramming in the pre-implantation embryo, it is not clear whether ARTs influence the integrity and stable transmission of the underlying DNA sequence itself. Prior investigations exploring this possibility have reached contradictory findings. Some studies have concluded that ARTs are associated with an increased burden of structural mutations and aneuploidy (29–31), while others have reported no significant increase in large-scale structural rearrangements in ART-compared to naturally-conceived offspring (32). Similarly, whereas a prior study in mice found no evidence for an increased point mutation rate in animals born via in vitro fertilization (IVF) (33), a recent retrospective comparative analysis in humans uncovered a significantly higher de novo point mutation rate in IVF-born children (34). However, the underlying etiology of sub- or in-fertility may predispose patients seeking ART to elevated rates of germline genome instability (35), such that any observed increase in the mutation burden associated with ARTs is potentially driven by ascertainment bias.

Despite conflicting earlier findings, there are multiple compelling lines of evidence to support the hypothesis that ARTs may be intrinsically mutagenic. First, local epigenetic features, including methylation at CpG dinucleotides and H3K9me3 marks, directly shape local mutation rates (36, 37). Thus, the genome-wide epigenetic dysregulation observed in ART-derived embryos could secondarily alter the genomic distribution and rate of new mutations compared to natural conceptuses. Second, the first few early embryonic cell divisions are especially error-prone and uniquely vulnerable to the accumulation of large-scale structural mutations and aneuploidies (38, 39). Exposure to stressors unique to the *in vitro* environment may exacerbate genomic instability at this critical time point in early development. For example, ART-derived embryos are often cultured at oxygen levels that do not precisely replicate those encountered in the maternal reproductive tract (40). Importantly, exposure to either hyperoxic or hypoxic culture conditions can elicit DNA damage and promote genomic instability (41, 42). Additionally, compounds such as glucocorticoids present in culture media could also perturb the early epigenetic landscape of the developing embryo, leading to widespread regulatory and cellular metabolic changes with downstream implications for genome integrity (43).

Here, we harness the efficiency of assisted reproductive technologies in the laboratory mouse to rigorously test whether a standard regimen of ARTs consisting of (1) ovarian hyperstimulation via administration of exogenous hormones, (2) gamete isolation and culture, (3) *in vitro* fertilization (IVF), (4) embryo culture, and (5) embryo transplantation into a receptive uterus influences the stable transmission of the mammalian genome. We use whole genome sequencing to identify *de novo* mutations in cohorts of age-matched ART-derived and natural-born mice. Our reliance on a single inbred strain mouse model allows us to rigorously control for potential genetic background effects and environmental exposures on mutation rates, eliminating these confounding factors from our analysis and overcoming a major limitation of retrospective clinical studies. We find a significant increase in the rate of *de novo* single nucleotide variants (dnSNVs) in ART-born mice, suggesting that this standard ART protocol adversely impacts genome integrity.

## RESULTS

### De novo single nucleotide variant discovery in naturally- and ART-conceived mice

We aimed to test whether a standard ART protocol affects the overall incidence of dnSNVs accrued in gametes and/or during early embryonic development in a mouse model. Specifically, we focus on the sequential procedures of ovarian hyperstimulation by exogenous hormone injection, oocyte collection, sperm isolation, gamete culture, *in vitro* fertilization, embryo culture to the 2-cell stage (∼1 day post fertilization), and embryo transfer to a pseudo-pregnant host dam. This series of ART procedures is routinely employed in the course of mouse husbandry for strain rederivation and colony expansion, and parallels protocols used in human fertility clinics with some exceptions. Notably, human embryos are typically cultured to the blastocyst stage (∼5 days post fertilization), and in most cases, undergo a freeze-thaw cycle prior to transfer.

We generated natural- and ART-derived cohorts of age-matched C57BL/6J mice derived from a common G0 founder pair (**Figure 1a**). Two G1 females were mated to a single male sibling, yielding two litters of naturally conceived mice. To obtain a matched ART-derived cohort, oocytes from two hormonally super-ovulated G1 females were harvested and *in vitro* fertilized with sperm from a second G1 male littermate. Sixteen 2-cell embryos derived from each of the two females were then transferred to the uteri of two pseudo-pregnant C57BL/6J recipient dams and reared to term. A total of 16 naturally born and 12 ART-conceived G2 progeny were whole genome sequenced to ∼50-72x coverage using two independent sequencing libraries (**Supplemental Table 1; Supplementary Figure 1**). The six G1 parents and two G0 pedigree founders were additionally sequenced to enable discrimination between dnSNVs and segregating variants within our pedigree (>75x coverage; 2 libraries per sample; **Supplemental Table 1; Supplemental Figure 2**). By bottlenecking the breeders used in this experiment through a single G0 founder pair and using a common sire for each treatment group, we minimize the number of background segregating mutations present in our G2 cohorts, limiting this potential source of false positive dnSNVs. Simulations based on empirical coverage estimates indicate that our study design has 80% power to detect a ∼30% increase in the dnSNV rate between the ART- and naturally-conceived cohorts, assuming a baseline mutation rate equivalent to that previously reported for C57BL/6J mice (46) (**Supplemental Figure 3**).

**Figure 1.**
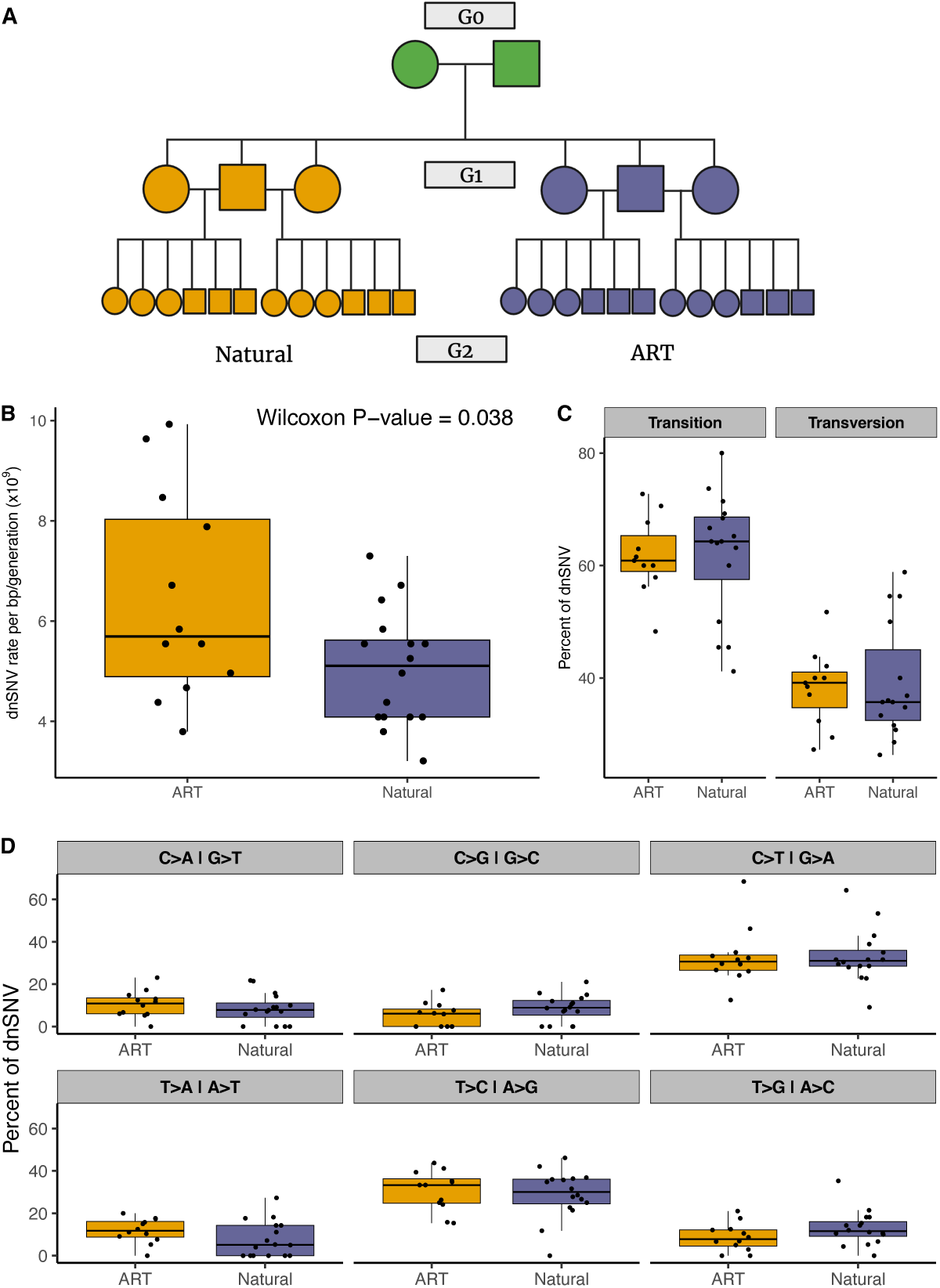
(A) A single founder mating pair (green; G0) produced six G1 breeders that were used to initialize two cohorts, one derived by natural mating (purple) and the other derived through a series of ARTs (orange). Each cohort consists of two litters of half-siblings. Circles represent females and squares represent males. (B) Boxplots showing the median, interquartile range, and full range of dnSNV rates per bp/generation across G2 samples. (C) Boxplots summarizing sample- and cohort-level variation in the relative transition and transversion rates and (D) mutational types.

Sequencing reads from each sample were aligned to the GRCm39 mouse reference genome, followed by SNV calling, dnSNV identification, and application of a post hoc filtering pipeline informed by tailored simulations (44) (see **Methods**). Our strategic use of the reference strain, C57BL/6J, effectively eliminates reference genome biases to maximize the accuracy of SNV calling and dnSNV discovery. On average, we identified 17 high-confidence autosomal dnSNVs in samples from the natural mating cohort (range = 11-25), corresponding to an estimated per base mutation rate of 3.88 x 10^-9^ per generation (per sample range: 2.47×10^−9^ to 5.61 x 10^-9^). This mutation rate estimate matches previously reported mutation rates for house mice (45–48). In contrast, offspring generated via ART exhibit a significantly elevated dnSNV burden, with an average of 22 autosomal dnSNVs per individual and an average dnSNV mutation rate of 4.95 x 10^-9^ (range: 13–29; range of mutation rate per sample: 2.91 x 10^−9^ to 6.50 x 10^−9^; one-tailed Wilcoxon rank-sum test, P-value = 0.038; **Figure 1B**).

Our finding of a significant increase in dnSNVs in ART-derived samples suggests that the tested ART protocol has adverse impacts on genome stability in gametes, zygotes, or early-stage embryos. However, systematic differences in sequencing data quality could lead to differences in the number of false positive (or false negative) dnSNV calls between cohorts. Assuredly, sequencing data quality is universally high across our G2 samples (**Supplementary Figure 1**), dnSNV calls are supported by similar variant quality metrics in both cohorts (**Supplementary Figure 4**), and our findings are recapitulated using sequencing data from individual replicate libraries from each sample (see Methods; **Supplementary Tables 3 and 4; Supplementary Figure 5**). We conclude that the observed differences in mutation burden between ART- and naturally-conceived offspring are not attributable to cohort-specific technical artifacts.

### No difference in mutation spectra between naturally conceived and ART-born mice

Exposure to specific environmental mutagens can impact the spectrum of mutations that accumulate in somatic tissues (49, 50), leading us to wonder whether the altered hormonal milieu and *in vitro* environment encountered during the generation of ART-born mice modifies the prevalence of specific types of mutations compared to those recovered in naturally-conceived animals. Both breeding cohorts exhibit qualitatively similar transition and transversion fractions (Wilcoxon rank sum, P-value > 0.05; **Figure 1C**), with estimates closely approximating those from prior studies of germline mutations in house mice (46, 51). Furthermore, the two cohorts show no difference in the relative frequency of individual mutation types (One-sided Wilcoxon rank sum test, P > 0.05) or the overall mutation spectrum (G-test, P-value > 0.05; **Figure 1D**). Further partitioning of dnSNVs based on their flanking nucleotide contexts reveals a significant increase in the C[C>A]A trinucleotide mutation fraction in ART-compared to naturally born samples (modified Chi-square *P*-value=0.019, **Figure 2c**). However, we note that resulting trinucleotide mutation count matrix is sparse, and we find no difference in the proportions of dnSNVs ascribable to defined mutational signatures between our two cohorts (**Figure 2b**). Overall, these findings reveal broad similarities in the mutation spectrum between ART- and natural-born mice, with the caveat that our analysis is likely underpowered to find differences.

**Figure 2.**
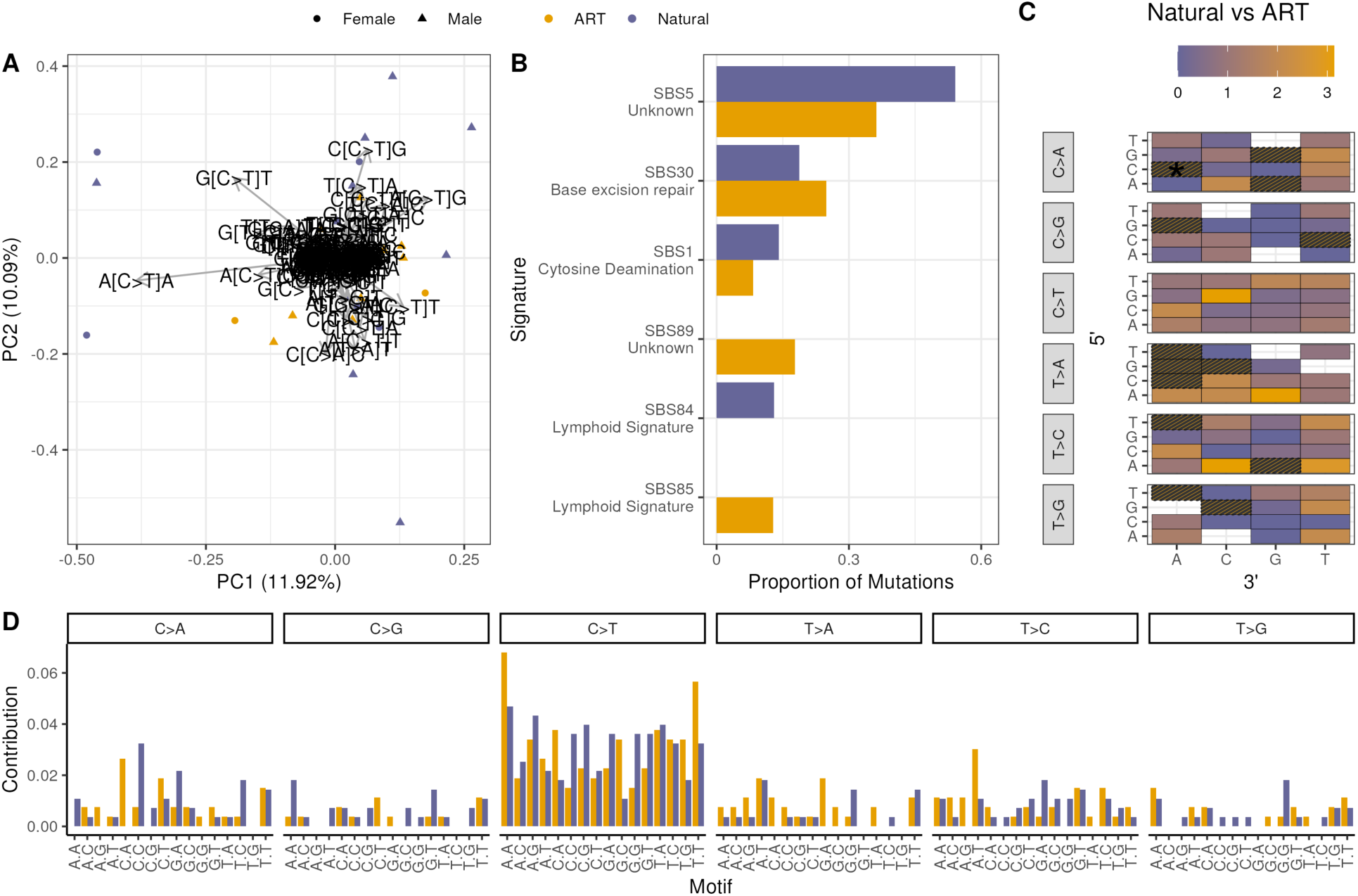
Comparison of mutational trinucleotide context and COSMIC single base signatures across ART and natural-born G2 samples. (A) Principal component analysis of the relative proportions of all 96 possible trinucleotide mutation types in the two birth cohorts. PC loadings (gray arrows) are plotted and labeled by the influential trinucleotide mutation. Points indicate the position of each sample along PC1 and PC2, with shapes representing sex and color denoting cohort. (B) Bar plot of the relative contribution of COSMIC single base signatures to the proportion of observed dnSNVs in the ART and natural birth cohorts. Permutation testing indicates no significant differences in signature contributions between cohorts. (C) Heatmap displaying the ratio of the average proportions of each of the 96 trinucleotide mutational contexts between ART and natural samples. White boxes indicate that the mutational type was absent in both cohorts, and grey hashing indicates that the mutation type was only present in ART samples and absent in Natural, rendering ratios as undefined. The asterisk indicates a significant difference in the C[C>A]A trinucleotide mutation fraction between the two cohorts. (D) Mutation spectra plot of the abundance of each of the 96 trinucleotide motifs in ART and natural-born samples.

### Evaluating the genomic landscape of dnSNVs in Natural- and ART-born mice

We next sought to understand whether the distribution of new mutations differs between our two cohorts with respect to various genomic features, including GC content, functional annotations, and repetitive elements. dnSNVs ascertained in both cohorts are uniformly distributed across the genome (Poisson test applied to mutation counts in 10Mb bins, *P* > 0.3; **Supplementary Figure 6A**) and arise in regions of similar GC-content (Wilcoxon rank sum, *P* = 0.892, **Supplementary Figure 6B**). Likewise, we detect no significant enrichment or depletion of ART-associated dnSNVs in CpG islands (Fisher’s exact test, P = 0.6862; **Supplementary Figure 6C**). The majority of dnSNVs occur in intronic and distal intergenic regions, with no cohort differences in the distribution of dnSNVs across different genomic annotations (Wilcoxon rank sum, *P* > 0.05; **Figure 3**). Similarly, there is no difference in the predicted variant effects of new mutations between natural- and ART-born mice (**Figure 3A**). The number of dnSNVs in ART-derived mice is modestly increased within LINEs, but the difference does not reach statistical significance (Chi-square Test, Bonferroni-corrected *P* > 0.13; **Figure 3D**). Despite an overall increase in dnSNV rate in ART-born mice, the genomic landscape of dnSNVs is largely indistinguishable between breeding cohorts.

**Figure 3.**
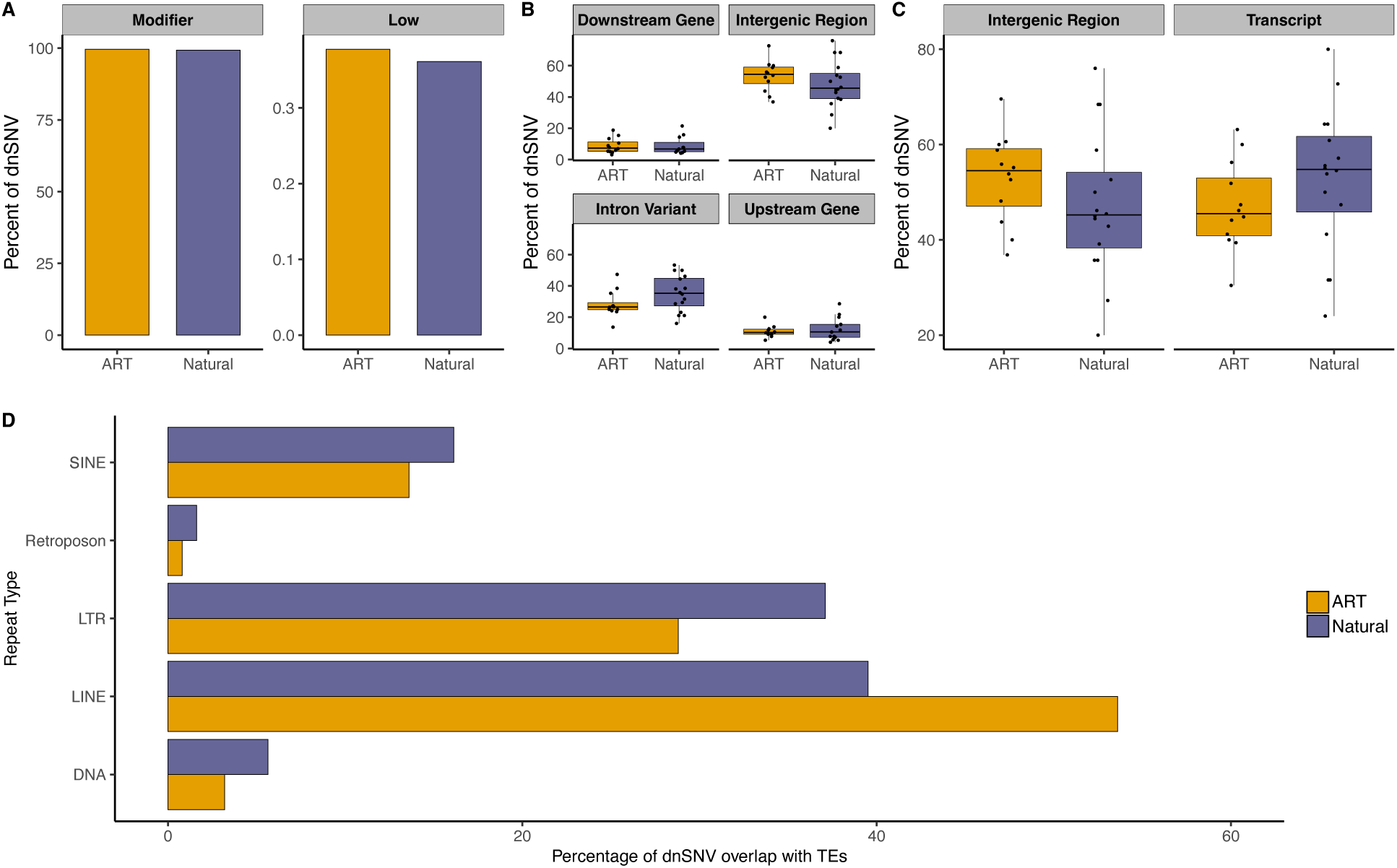
The genomic landscape of dnSNVs in naturally-born and ART-derived mice. (A) Proportion of dnSNVs in natural and ART genomes assigned to “Modifier” and “Low” functional effect predictions. Proportion of dnSNVs in that reside in (B) genomic regions assigned to various functional annotations and (C) within annotated transcripts versus intergenic regions. (D) The percentage of dnSNVs overlapping TE repeat classes in natural and ART-born mice. We find no significant cohort-level differences in dnSNV enrichment in any of these annotation categories.

### Contrasting the epigenomic and genome regulatory context of dnSNVs

Early development is characterized by the coordinated erasure, deposition, and redistribution of numerous DNA and histone modifications (52). Prior work has established that this epigenetic reprogramming is perturbed *in vitro* relative to *in vivo* (16). Given that diverse chromatin modifications are associated with intragenomic mutation rate variation (36), we next set out to explore whether dnSNVs arise in distinct epigenetic contexts in our two cohorts. We intersected the positions of dnSNVs ascertained in both the ART- and natural-born samples with genome-wide maps of several epigenetic marks in mouse embryonic stem cells (mESCs) (53) and early mouse embryos (54). There is no cohort-level difference in the proportion of dnSNVs that arise in regions associated with H3K27ac, H3K36me3, H3K4me1, H3K4me3, H3K9ac, H3K9me3 histone modifications in either mESCs or mouse embryos (Chi-square *P* > 0.4; **Supplementary Figures 7 and 8**).

In somatic cells, transcription-coupled repair processes lead to reduced dnSNV rates in highly expressed genes relative to more lowly expressed or transcriptionally silenced genes (55). Although differences in gene expression have been documented between ART- and naturally born progeny (17), we find no cohort-level difference in the transcriptional activity of genes harboring dnSNVs in C57BL/6J-derived mESCs (Wilcoxon Signed Rank Test, *P* > 0.2; **Supplementary Figure 9**).

A majority of dnSNVs are presumed to arise due to errors in DNA replication, with later-replicating regions of the genome exhibiting elevated mutation rates compared to early-replicating regions (56). Using published replication timing estimates derived from mESCs, we again find no evidence for differences in overall replication timing at dnSNV sites in our ART- and naturally-born cohorts (Mann Whitney U-Test, *P* > 0.05; **Supplementary Figure 10**) (57, 58).

### No difference in the developmental timing of dnSNVs between ART- and naturally-conceived mice

The excess dnSNVs recovered in our ART-born mice may have arisen during the (i) formation of gametes *in vivo*, (ii) *in vitro* gamete manipulation and culture, (iii) the transition from the zygote to 2-cell stage, or (iv) at later stages of embryonic development. We profiled the allele depth ratio (i.e., the proportion of sequencing reads supporting the alternative allele) across all dnSNVs to gain insight into the developmental timing of mutations in our two cohorts. dnSNVs that arose in parental gametes will be constitutively present in the genome of offspring and manifest as an allele depth ratio close to ∼0.5, whereas mutations that arise during early development will have lower allele depth ratios. We observed no significant difference in the allele depth ratios of dnSNVs between animals conceived via ART and those conceived naturally (Wilcoxon rank sum test, *P* = 0.795; **Supplementary Figure 11**), indicating no detectable difference in the temporal origin of dnSNVs between cohorts.

### No detectable difference in de novo structural variation rates between ART- and natural-born mice

A recent report identified a significant increase in the rate of de novo structural variants (SV) in cattle born via ART compared to naturally conceived animals (31). We utilized an ensemble SV calling approach to identify de novo deletions and duplications >50 bp in our G2 samples (see **Methods**). Eight and seven high-confidence germline *de novo* SVs were identified in the natural (4 deletions and 4 duplications) and ART-born samples (4 deletions and 3 duplications), respectively, an insignificant difference (one-tailed Wilcoxon rank sum test; *P* = 0.931; **Supplemental Table 5**). The rate of de novo SV is predicted to be at least one order of magnitude lower than the single nucleotide mutation rate (59), a consideration that renders our study insufficiently powered to find small or modest differences in SV mutation rate.

## DISCUSSION

Prior work has shown that ARTs are associated with epigenetic and transcriptomic changes in early embryos, but the potential impact of these procedures on DNA sequence integrity is not well-understood. Here, we took advantage of a well-controlled mouse model system to directly estimate the burden of dnSNVs in C57BL/6J inbred mouse cohorts conceived through a standardized ART protocol or via natural mating. We document a statistically significant increase in the overall dnSNV rate in ART-derived mice compared to their age- and genetically-matched naturally-born counterparts. Our dnSNV discovery pipeline surpasses field standards for rigor, relying on two independent sequencing libraries for each sample and invoking a simulation-informed protocol for dnSNV discovery in inbred mouse genomes (44).

The biological mechanisms underlying the elevated rate of dnSNVs in ART-derived mice remain unclear. Known mutational processes often leave distinct genomic signatures or show regional enrichment (50, 60), yet we found no such signals in our data. While the modest number of dnSNVs identified may obscure subtle differences in the mutational landscape, we find no differences in the transition/transversion ratio, the mutation spectrum, or the genomic distribution of dnSNVs between cohorts. Similarly, we find no differential enrichment for dnSNVs across various epigenetic contexts, or with respect to GC content, local transcriptional activity, or replication timing.

Future studies will be necessary to determine whether the elevated mutation burden observed in ART-derived mice arises from a specific step within the ART pipeline or reflects the cumulative effects of multiple procedures. One plausible contributor is ovarian stimulation using exogenous follicle-stimulating hormone (FSH) and human chorionic gonadotropin (HCG). These hormones induce the resumption of meiosis in oocytes, a process known to be highly error-prone (61). Hormone-induced ovulation could alter the fidelity of meiotic double-strand break repair or impair chromosome segregation, thereby contributing to the increased rate of *de novo* mutation. Although speculative, this possibility has some support. Ovulation induction has been associated with elevated risks of miscarriage and congenital anomalies in humans (62, 63), and a recent retrospective analysis suggested a link between this intervention and increased maternally transmitted mutations (34). Nonetheless, we cannot exclude the potential influence of other ART-related factors, such as mechanical stress during embryo manipulation or the physicochemical properties of the *in vitro* culture environment. Our use of male and female parents from a single inbred strain limits the ability to assign the parental origin of the dnSNVs reported here, an advantage that could help localize the ultimate source for the excess dnSNV burden associated with ART.

Overall, the ∼30% mutation rate increase observed in ART-derived offspring is unlikely to have substantial implications for the mutation load in laboratory populations maintained via recurrent cycles of IVF-based rederivation (64) or husbandry programs that utilize ARTs to rederive strains on regular intervals to curtail genetic drift (65). Assuming that ∼0.5% of new single nucleotide mutations are strongly deleterious (66), a baseline mouse mutation rate of μ = 0.5 × 10^-9^/bp/gen (45, 46, 48), and genome size of 2.7Gb, we expect ∼0.067 new deleterious mutations per mouse per generation under a traditional breeding program (0.5 × 10^-9^ × 2.7Gb × 0.005 = 0.067). Using ART, the number of expected new deleterious mutations is increased to ∼0.088 per genome per generation. Thus, for every ∼50 ART-derived mice, one additional highly deleterious dnSNV is expected to be recovered compared to the natural mating baseline. The magnitude of this impact is roughly equivalent to an increase in mouse paternal age of ∼30 weeks (46).

While our findings reveal a moderate mutagenic impact of ARTs in mice, we emphasize that our conclusions cannot be readily extrapolated beyond the mouse model system profiled here. Notably, mice and humans differ in their reproductive physiology, dynamics of epigenetic reprogramming, and timeline of early embryonic development, considerations that may influence both the rate and spectrum of dnSNVs and the sensitivity of mutation accumulation under ART. Further, ART protocols employed in human fertility clinics often include prolonged embryo culture to the blastocyst stage and an embryo freeze-thaw cycle, steps that present clear departures from the use of fresh 2-cell stage transfers in our mouse protocol. Whether or how these protocol differences impact potential mutation rate differences associated with ART is unknown. While we caution that our work has no immediate implications for human clinical practice, our findings nonetheless strongly motivate further investigation to assess the potential mutagenic impact of ART in humans. This need is particularly urgent in view of forecasted trends of increasing reliance on ART due to sociodemographic shifts and increasing democratization of access to ARTs (5).

## MATERIALS AND METHODS

### Animal husbandry and establishment of breeding cohorts

A single C57BL/6J breeding pair was obtained from The Jackson Laboratory and housed in a low barrier room in accordance with an animal care protocol approved by The Jackson Laboratory’s Animal Care and Use Committee (Protocol #17021). Two G1 females from the initial litter born to this G0 C57BL/6J founder breeding pair were naturally mated to a single male G1 littermate. One of these mated G1 females produced a litter of 9 live-born pups and the second gave birth to a litter of 7 mice (**Figure 1a**). All 16 naturally born G2 pups were reared to 4 weeks of age and euthanized by exposure to CO_2_ prior to terminal tissue collection.

Two additional females and one male from the same G1 litter were transferred to The Jackson Laboratory’s Reproductive Services Facility at 4 weeks of age. Females were injected with 5 IU PMSG followed 48 hours later by a 5 IU HCG trigger to induce ovulation. Oocyte clutches were then harvested from the ampullae of each super-ovulated female and incubated in 150uL Cook RVF media supplemented with an additional 50uL of reduced glutathione (GSH) media at 37°C under mixed gas (5% CO_2_, 5% O_2_, and 90% N_2_) for 30-60 minutes. Concurrently, sperm were isolated from the caudal epididymides of the donor male and incubated in TYH media supplemented with 0.75mM Methyl-β-cyclodextrin at 37°C under mixed gas for 40-60 minutes.

Following incubation, egg clutches and 10uL of sperm were transferred to a 1mL drop of Cook RVF media covered with mineral oil. Fertilization was allowed to occur over a ∼2-6-hour period at 37°C under mixed gas. Zygotes were then washed by transfer through two sequential droplets of Cook RVF Media and incubated overnight in a final droplet of Cook RVF media at 37°C under mixed gas. Approximately 16 two-cell embryos from each donor female were subsequently transferred to the uteri of two pseudo-pregnant C57BL/6J dams and reared to term. A total of 29 live-born ART-derived G1 pups were euthanized by CO_2_ at approximately 4 weeks of age for terminal tissue harvests.

### DNA extraction, library preparation, and sequencing

The G1 founder breeding pair, 6 G1 parents, 16 naturally born G2 mice, and 12 ART-derived G2 mice were selected for whole-genome sequencing (n = 36 samples; **Figure 1a**). Genomic DNA isolation, library preparation and sequencing were performed in duplicate for all samples, including an initial round of data collection performed in 2020 and a second batch completed in 2023. Genomic DNA was isolated from snap frozen mouse tails using the NucleoMag Tissue Kit (Machery-Nagel) according to the manufacturer’s protocol. DNA concentration and quality were assessed using the Nanodrop 8000 spectrophotometer (Thermo Scientific), the Qubit Flex dsDNA BR Assay (Thermo Scientific), and the Genomic DNA ScreenTape Analysis Assay (Agilent Technologies) and judged to be sufficient for library preparation for all samples. Paired-end 150bp whole genome libraries were constructed using the KAPA HyperPrep Kit (Roche Sequencing and Life Science) according to the manufacturer’s protocols, targeting an insert size of 400 base pairs. Briefly, the protocol entails shearing the DNA using the E220 Focused-ultrasonicator (Covaris), size selection targeting 400 bp, ligation of Illumina specific barcoded adapters and 9bp UMI adaptors, and 1 cycle of PCR amplification. Library quality was assessed using the D5000 ScreenTape (Agilent Technologies) and concentration was determined by a Qubit dsDNA HS Assay (ThermoFisher).

The initial set of paired-end 150bp Illumina libraries were prepared and sequenced to ∼20-30x coverage on an Illumina NovaSeq6000 using a combination of S2 and S4 flow cells. A second set of paired-end 150bp libraries were pooled and sequenced to ∼30x coverage (∼90 Gb/sample) on an Illumina NovaSeq 6000 using the S4 Reagent Kit v1.5. For clarity, we refer to sequencing data from these two libraries as “Seq1” and “Seq2”. All sequencing data are available on the NCBI Sequence Read Archive under PRJNA1282662.

### Read processing and mapping

Read quality assessment and adaptor trimming were performed on each sample library using fastp (v. 0.23.4) (67). Processed reads were then mapped to GRCm39 mouse reference genome using default parameters in *bwa mem* (v. 0.7.17-r1188), indexed, and processed for duplicate read discovery using *samtools* (v. 1.21). Duplicate reads were identified by executing the following piped command series:

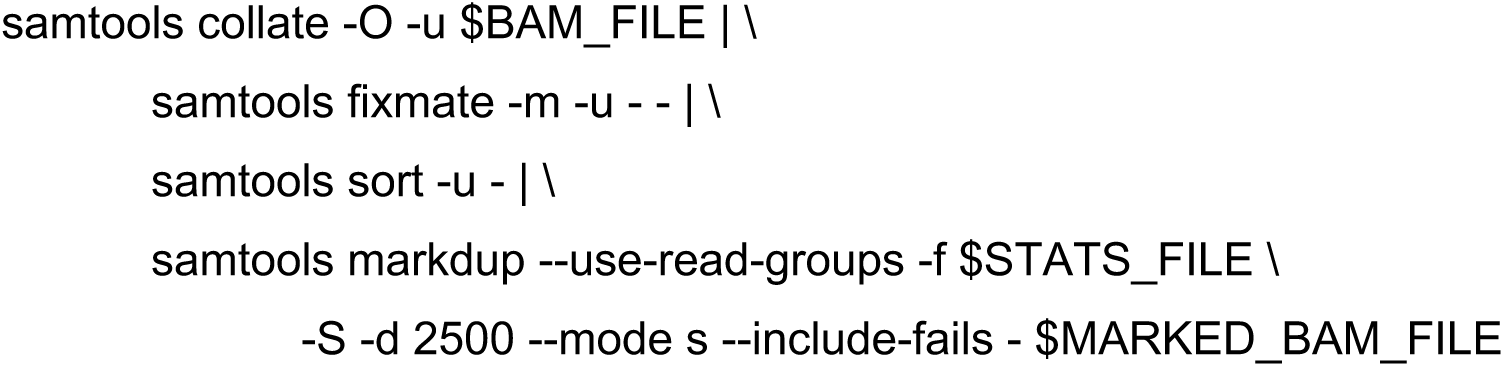

Mapping metrics were computed for each sequenced library by invoking the *flagstat* and *idxstats* command in samtools (**Supplementary Table 1**). The two independent libraries generated from each sample were then merged using samtools merge, and mapping metrics re-computed on the merged file (**Supplementary Table 1**).

### Single nucleotide variant calling

Single sample variant calling was performed using DeepVariant (v. 1.6.1), invoking the pre-trained WGS model (68). A joint callset including all 36 samples was derived using GLnexus (v. 1.2.7). In parallel, we used Mpileup (via the ‘bcftools call’ command in bcftools v. 1.9-1) with default parameters to produce a second joint callset. These variant calling steps were executed on the individual Seq1 and Seq2 bam files from each sample, as well as the merged bam file integrating data from both Seq1 and Seq2. Calls from the merged data were used as the primary source for dnSNV discovery (see below), with the Seq1 and Seq2 call sets offering supportive confirmation of dnSNVs in two independent libraries.

### De novo mutation discovery

dnSNV discovery was performed using the joint SNV call sets generated from merged BAM files containing reads from both sequencing libraries (Seq1 and Seq2). The joint VCF file was filtered using bcftools (v1.16) to retain only autosomal, biallelic single nucleotide variants that were present in a heterozygous state in a single G2 individual and absent from all other individuals in the pedigree. Following our simulation-based recommendations for dnSNV discovery in mice, we retained only those calls supported by both Mpileup and DeepVariant and applied regional filters to eliminate dnSNVs residing in genomic regions prone to false positive calls (44). Specifically, putative dnSNVs were excluded if they (1) overlapped with any of the following annotations defined by the repeatMasker track accessed from the UCSC Genome Browser (69): low complexity regions, rRNA, satellite DNA, scRNA, simple repeats, snRNA, srpRNA, and tRNA; (2) were flanked on one or both sides by an A/T homopolymer run of at least 5 bp; (3) were located within 35 bp of another SNV; or (4) overlapped a manually curated set of highly copy number variable genes in the mouse genome (**Supplemental Table 6**). We next assured that putative G2 dnSNVs surviving these strict filters were independently supported by calls in both Seq1 and Seq2. The final set of dnSNVs is provided in **Supplemental Table 2**.

We employed an identical procedure for dnSNV discovery in the individual Seq1 and Seq2 data sets, excluding the requirement that dnSNVs are confirmed by both sequence batches (**Supplemental Tables 3 and 4**).

### De novo mutation rate estimation and analysis of mutation spectrum

The per base *de novo* mutation rate (μ) was estimated for each G2 sample using the following formula:

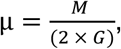

where *M* is the number of dnSNVs per G2 sample, and *G* is effective haploid genome size in base pairs. G was calculated as the total length of the 19 mouse autosomes (2.44 Gb) the minus masked regions described above, resulting in an effective genome size of 2.23 Gb. dnSNV spectra were obtained for each G2 sample using the TsTv-summary output flag in VCFtools version 0.1.16 (70). dnSNVs were further annotated by their trinucleotide context (i.e., the focal dnSNV and its immediate 3’ and 5’ flanking nucleotides) using the R libraries SomaticSignatures (version 2.38.2, Gehring et al. 2015) and VariantAnnotation (version 1.48.1, Obenchain et al. 2014). The proportion of mutations in a given sample that fall into each mutational category was used as input for a principal component analysis to evaluate variation in the dnSNV spectrum between ART- and natural-born mice. We used a modified Chi-square test with *P*-values corrected for non-independence to test for differences in the mutation fraction between the two cohorts. Finally, we aggregated the trinucleotide spectra across our two breeding cohorts to assess relationships with defined COSMIC single base mutation signatures (v 3.4) (50) computed for mm10 using SigProfilerAssignment in Python (version 0.2.3, (71)). We excluded mutation signatures not relevant to our unexposed cohort (e.g., Arisocholic Acid, Artifact, Colibactin, UV damage, Tobacco, Treatment Signatures, and immunosuppressants). To evaluate the statistical significance of signature contributions between the two groups, we randomly shuffled the cohort labels of the dnSNVs in their trinucleotide contexts and re-computed mutation signatures. We then compared the observed cohort difference in the proportion of each trinucleotide mutation class to the distribution of 1000 randomly simulated differences. A 1-sided *P*-value was calculated as the probability of observing a difference as large as or larger than the observed difference. All statistical analyses were performed using R (4.2.2) and RStudio (4.2.2).

### Genomic annotation and epigenomic enrichment of dnSNVs

dnSNVs were annotated using SnpEff (v. 5.0, (72)). *Bedtools intersect* (v2.28.0) was used to determine the numbers of dnSNVs overlapping various classes of repeat elements annotated in the mm39 references using the RepeatMasker track extracted from UCSC Table Browser (69). Similarly, we interrogated SNP locations over CpG Islands by intersecting dnSNVs with the ‘CpG Island’ track from the UCSC Genome Browser. To contrast the GC content of dnSNVs in the natural and ART-derived cohorts, we calculated the GC content of a 201bp window centered on each dnSNV and compared the distributions of flanking GC content.

dnSNVs were intersected with CTCF binding sites and various histone modifications assayed by ChIP-seq in C57BL/6J mouse ESCs under the mouse ENCODE project (73) and early mouse embryos (54). dnSNV coordinates were first lifted over to mm10 reference genome coordinates to ensure compatibility with ChIP-seq peaks associated with ENCODE datasets. Differential enrichment of dnSNVs between cohorts was assessed by Chi-Square tests. Similarly, dnSNVs were intersected with quantitative estimates of transcript abundance in C57BL/6J mouse ESCs (73), allowing a window of 2.5kb upstream and downstream of the gene start and end coordinates, respectively. Cohort differences in the mean expression level of genes neighboring dnSNVs and the proportion of dnSNVs neighboring active versus inactive genes were assessed by a Wilcoxon Rank Sum test and a Chi-square test, respectively. To evaluate potential cohort differences in the replication timing of genomic regions where dnSNVs arise, dnSNVs were intersected with published Repli-seq replication timing estimates on mESCs (57, 58). A Wilcoxon rank sum test was used to assess significance.

### Structural variant calling and de novo structural variant discovery

To identify potential *de novo* structural mutations, we performed SV discovery on each cohort using DELLY (v.0.8.7) (74) and Manta 1.6.0 (75). We used default settings in DELLY to perform per sample germline SV calling against the GRCm39 reference and subsequently merged calls across all samples in our pedigree. In parallel, we used Manta to jointly call germline SVs in each of the 28 parent-offspring trios embedded in our pedigree (**Figure 1A**), followed by merging of these per trio SV call sets. (Running a joint sample analysis with Manta on larger sample cohorts caused run time challenges and proved to be infeasible with our compute resources). We then intersected the two final SV call sets from Manta and DELLY using the collapse command in Truvari (v4.0.0) (76) with the following parameters specified: -pctsize 0.75 –pctovl 0.5 –pctseq 0.7-s 20 -S 10000000 -k common --chain. We retained only the calls that were supported by both callers and that were unique to a single G2 sample. We focus on deletions and duplications, to the exclusion of complex and copy number natural SVs, owing to the inherent limitations of short-read data. These candidate de novo structural variants were then visually inspected for read depth signatures consistent with duplications and deletion calls using Samplot (version 1.1.6; (77)). Only calls visually supported by expected read depth patterns were retained. This manual filter resulted in the exclusion of 462 deletions and 108 duplications. De novo SVs were annotated for predicted functional effects using the Ensembl variation effect predictor (VEP; tool version 2.0; (78)).

## Supporting information

simulation code for R

## ACKNOWLEDGEMENTS

We thank members of the Dumont Lab, Baker Lab, and Mary Ann Handel at The Jackson Laboratory for critical feedback on this project. We are indebted to the technical expertise of the scientific staff in The Jackson Laboratory’s Reproductive Sciences and Genome Technologies Scientific Service for carrying out ART procedures and whole genome sequencing, respectively. We also thank the Research IT Staff at The Jackson Laboratory for their oversight and maintenance of the high-performance computing resources that made this work possible. This work was supported by start-up funds from The Jackson Laboratory and a MIRA from The National Institute of General Medical Sciences to BLD (R35 GM133415).

## SUPPLEMENTARY FIGURES

**Supplemental Figure 1.**
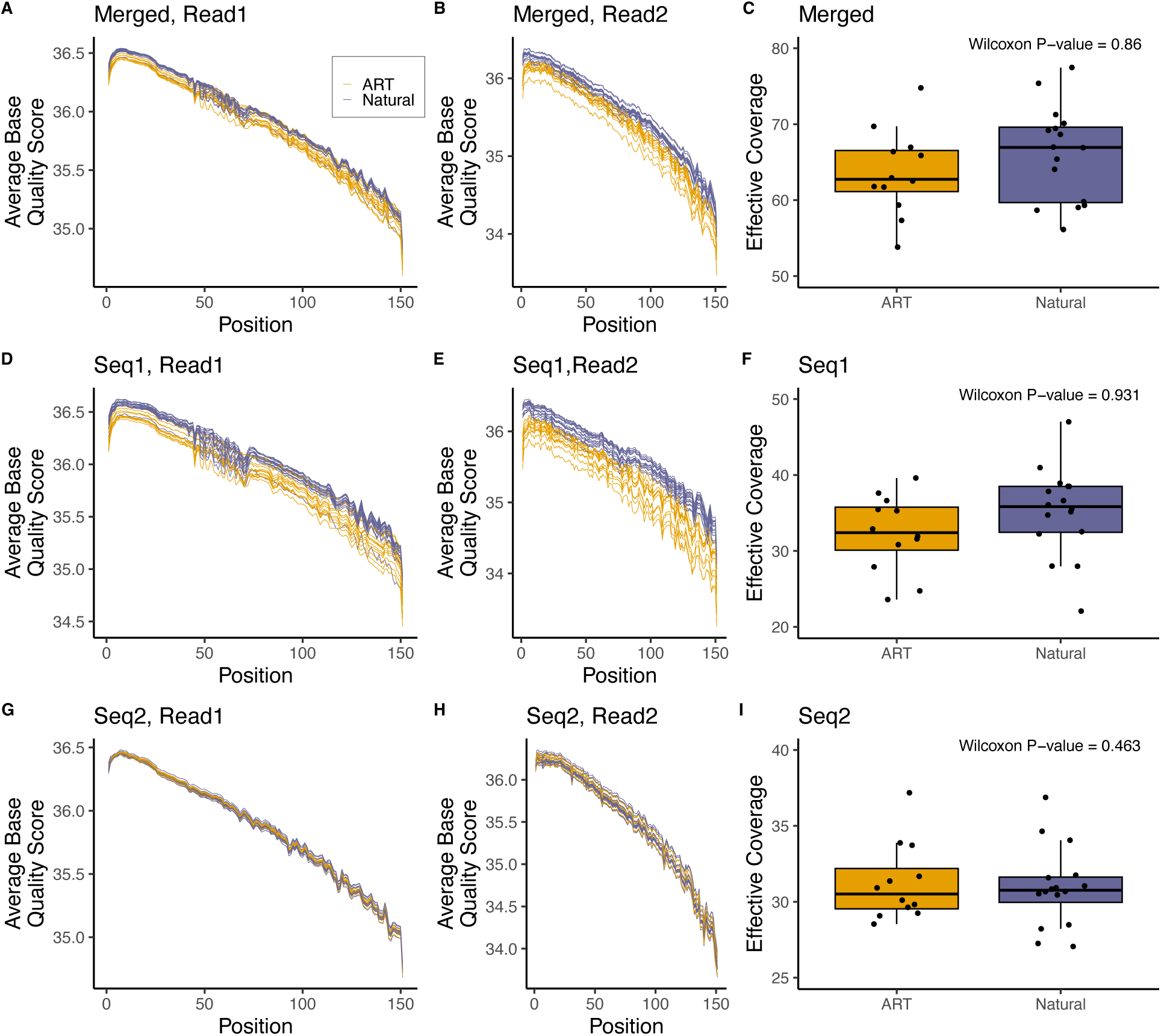
Base quality scores by position in read and sequencing coverage for ART-(orange) and natural-born samples (purple). Two replicate paired-end 150bp libraries were prepared from each sample and sequenced in two batches, referred to as Seq1 and Seq2. Average base quality scores across each base position in Read1 in the merged data (A) and both individual sequencing libraries (D, G). Average base quality scores across each base position in Read2 in the merged data (B), and individual Seq1 (E) and Seq2 (H) libraries. High-quality sequencing reads were obtained from both breeding cohorts in both sequencing batches, although reads from ART-derived samples have slightly lower base quality scores compared to natural-born samples in the Seq1 sequencing batch. Effective sequencing coverage for the merged, Seq1, and Seq2 data (C, F, I). Coverage estimates exclude duplicated reads and assume a genome size of 2.7Gb. Samples from both ART- and natural-born cohorts were sequenced to identical target coverage in both Seq1 and Seq2. However, a batch effect in sample preparation, library construction, and sequencing manifested as a higher rate of duplicated reads in the ART-derived samples compared to natural-born samples during Seq1 data collection (F).

**Supplemental Figure 2.**
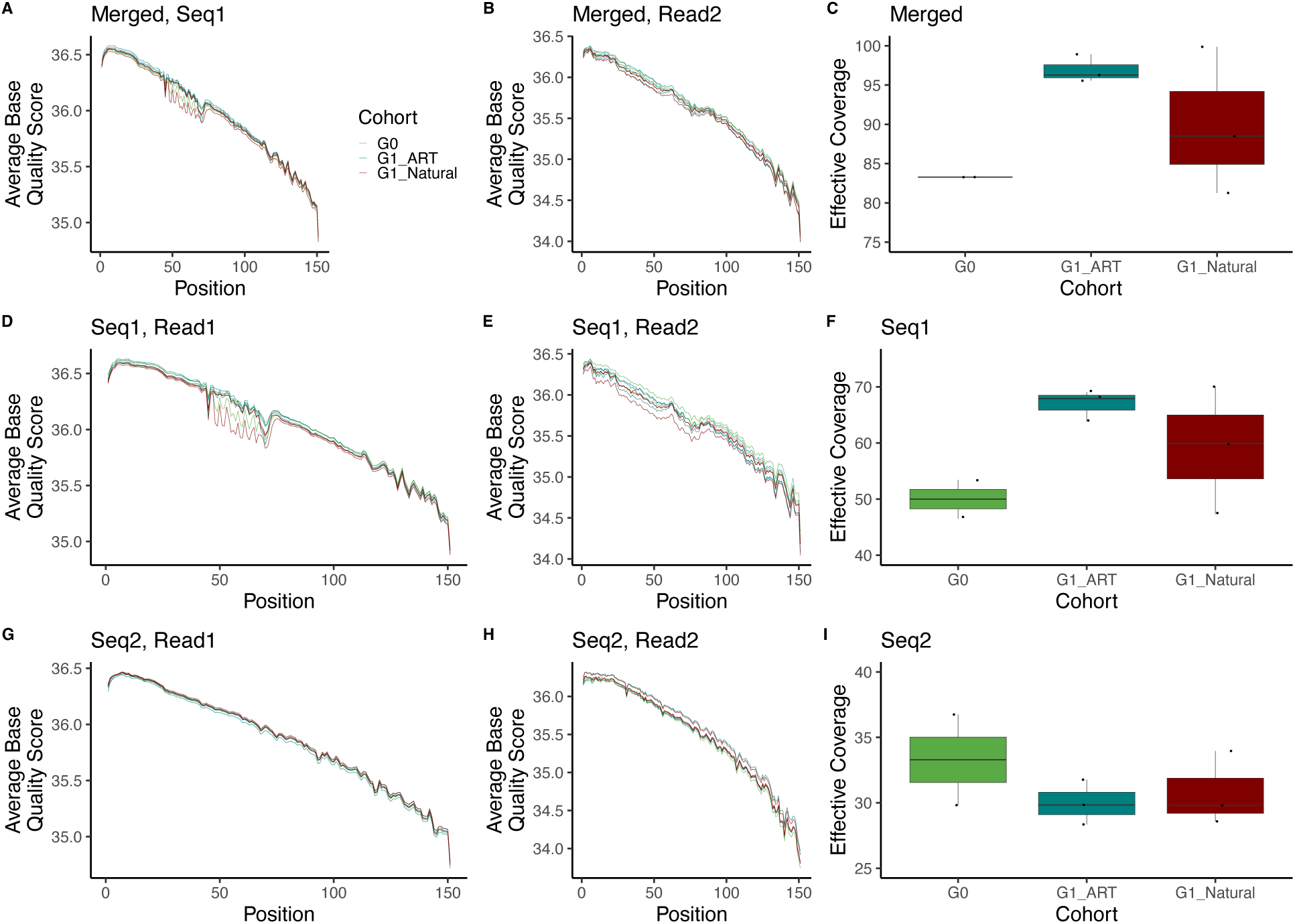
Base quality scores by position in read and sequencing coverage for the G0 pedigree founder pair and G1 parents of the ART- and natural-born cohorts. Two libraries were prepared from each sample and processed in two different sequencing runs, Seq1 and Seq2. Average base quality scores across each base position in Read1 for the merged sequencing data (A) and individual Seq1 (D) and Seq2 (G) sequence data. Average base quality scores across each base position in Read2 for the merged, Seq1, and Seq2 sequence data (B, E, H). Effective sequencing coverage for the merged, Seq1, and Seq2 sequence data (C, F, I).

**Supplementary Figure 3.**
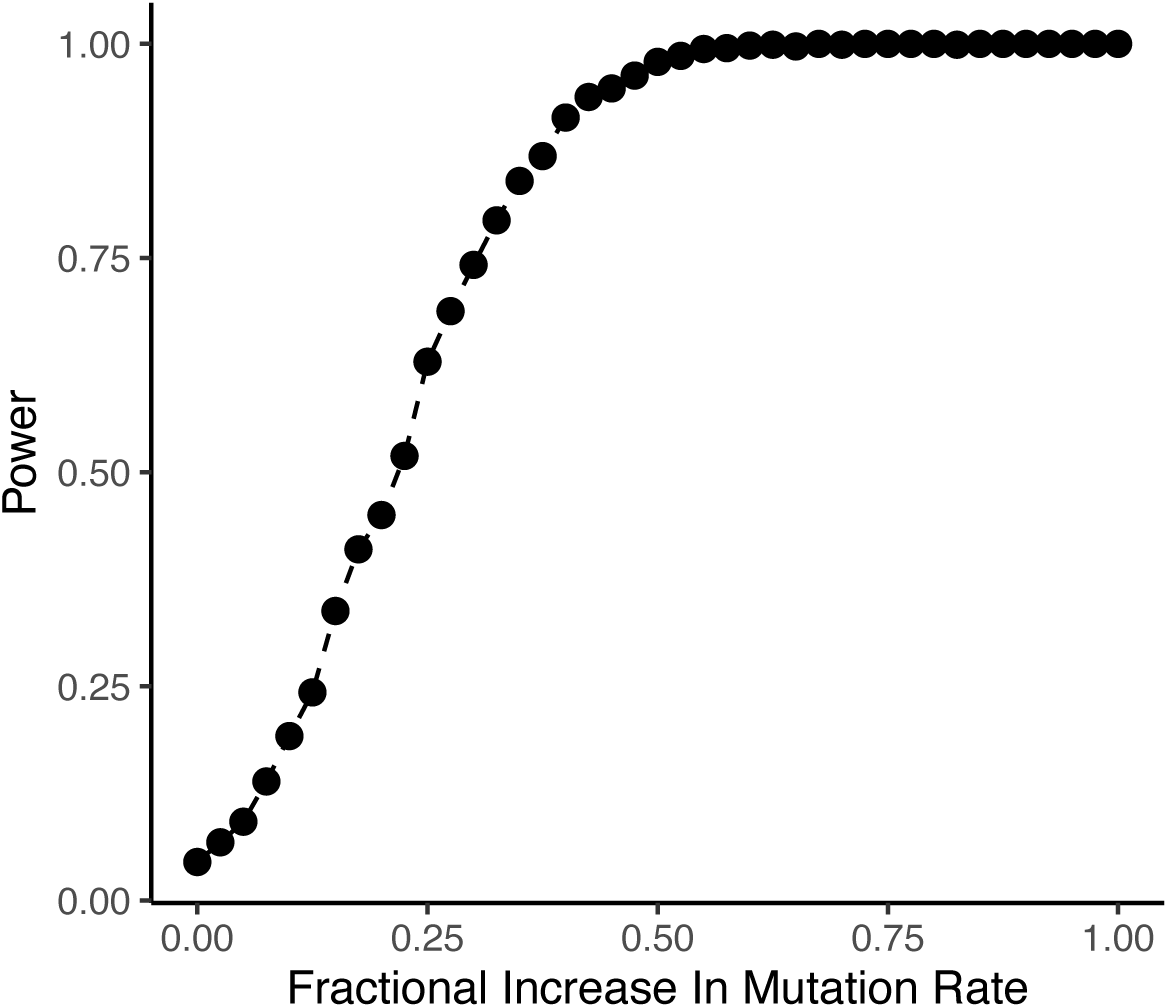
Power to detect mutation rate differences under the analyzed experimental conditions. We simulated dnSNVs in each G2 sample according to a Poisson distribution, with rate parameter set to the number of expected germline mutations (genome size x mutation rate in per base/per generation units). Simulations assume an average callable genome size of 2.23 Gb, a baseline mutation rate of 0.5 x 10 per base per generation, and impose a read depth requirements of ≥10 total reads and ≥3 reads per allele. Binomial sampling was used to capture the uncertainty associated with transmission of mutations arising in the G1 germline to the G2 generation. The mutation rate of the simulated natural cohort was held at the simulated baseline rate, whereas the mutation rate of the simulated IVF cohort was increased by a fixed proportion across simulation sets (x-axis). The difference in mutation tallies among the simulated natural and ART cohorts was assessed by a one-way Wilcoxon rank sum test. We performed a total of 1000 simulation replicates for each fractional increase in mutation rate in the ART cohort, retaining the Wilcoxon rank sum test P-value from each replicate. Power was computed as the fraction of simulated datasets for which P<0.05. R code to reproduce simulations and this figure is available as supplemental material.

**Supplementary Figure 4.**
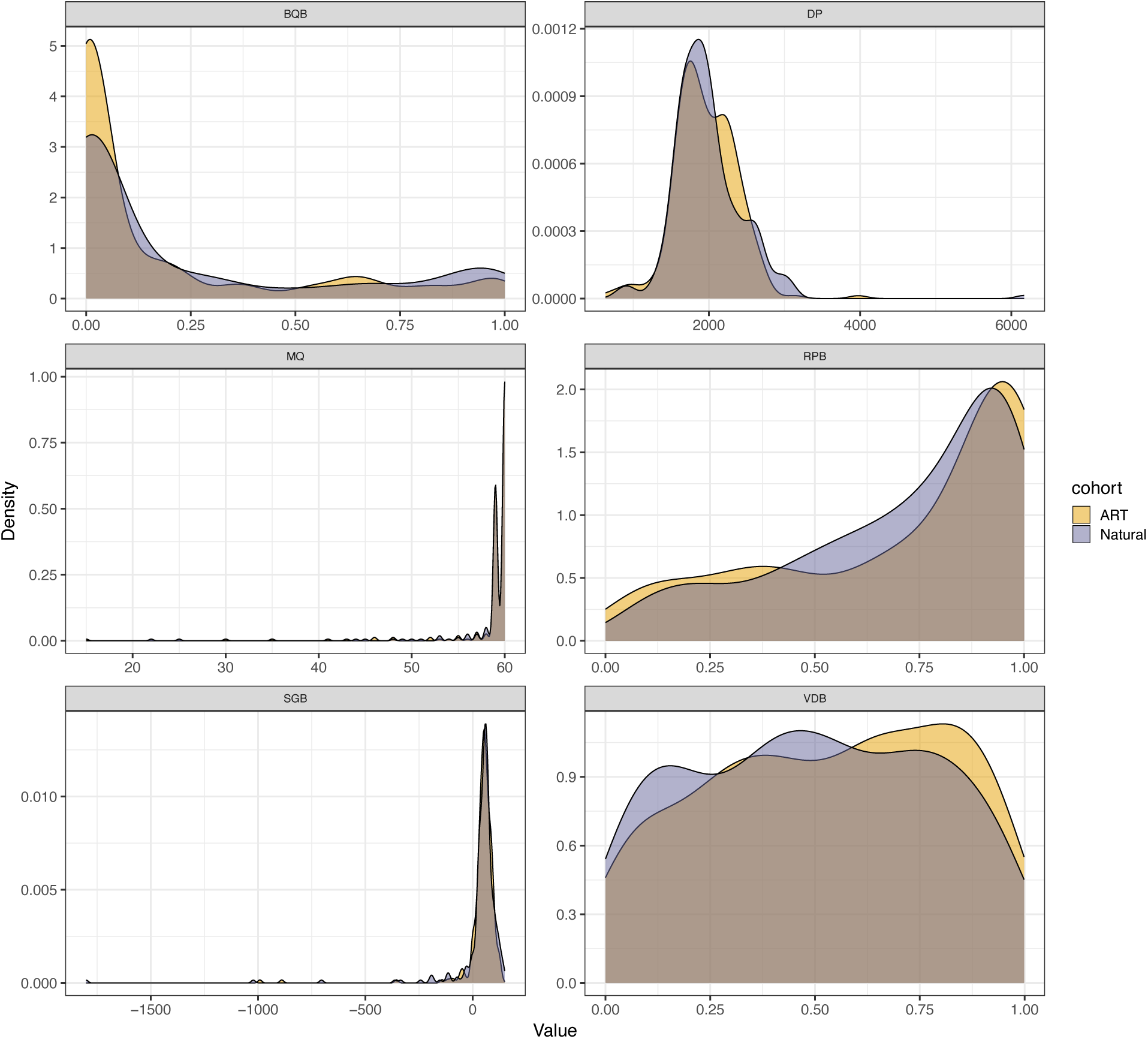
Distribution of Mpileup variant annotations for dnSNVs in ART-(yellow) and naturally-conceived (purple) progeny. Annotations include Base Quality Bias (BQB), Depth of Coverage (DP), Mapping Quality (MQ), Read Position Bias (RPB), Segregation-Based Score (SGB), and Variant Distance Bias (VDB). Annotation distributions are identical between the two groups (Kolmogorov-Smirnov Test, P > 0.05).

**Supplementary Figure 5.**
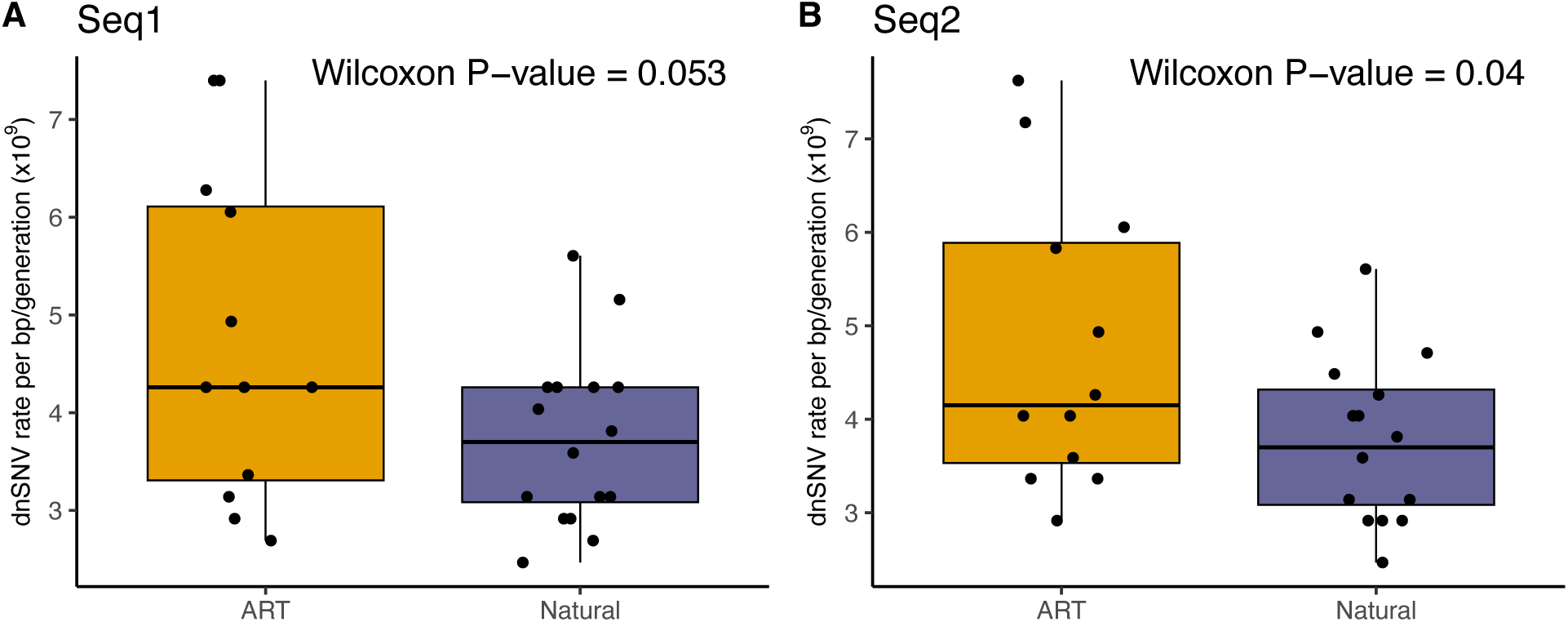
Boxplots showing the median, interquartile range, and full range of dnSNV rates per bp/generation across G2 samples. dnSNV calls were derived using only sequencing data from (A) Seq1 and (B) Seq2.

**Supplementary Figure 6.**
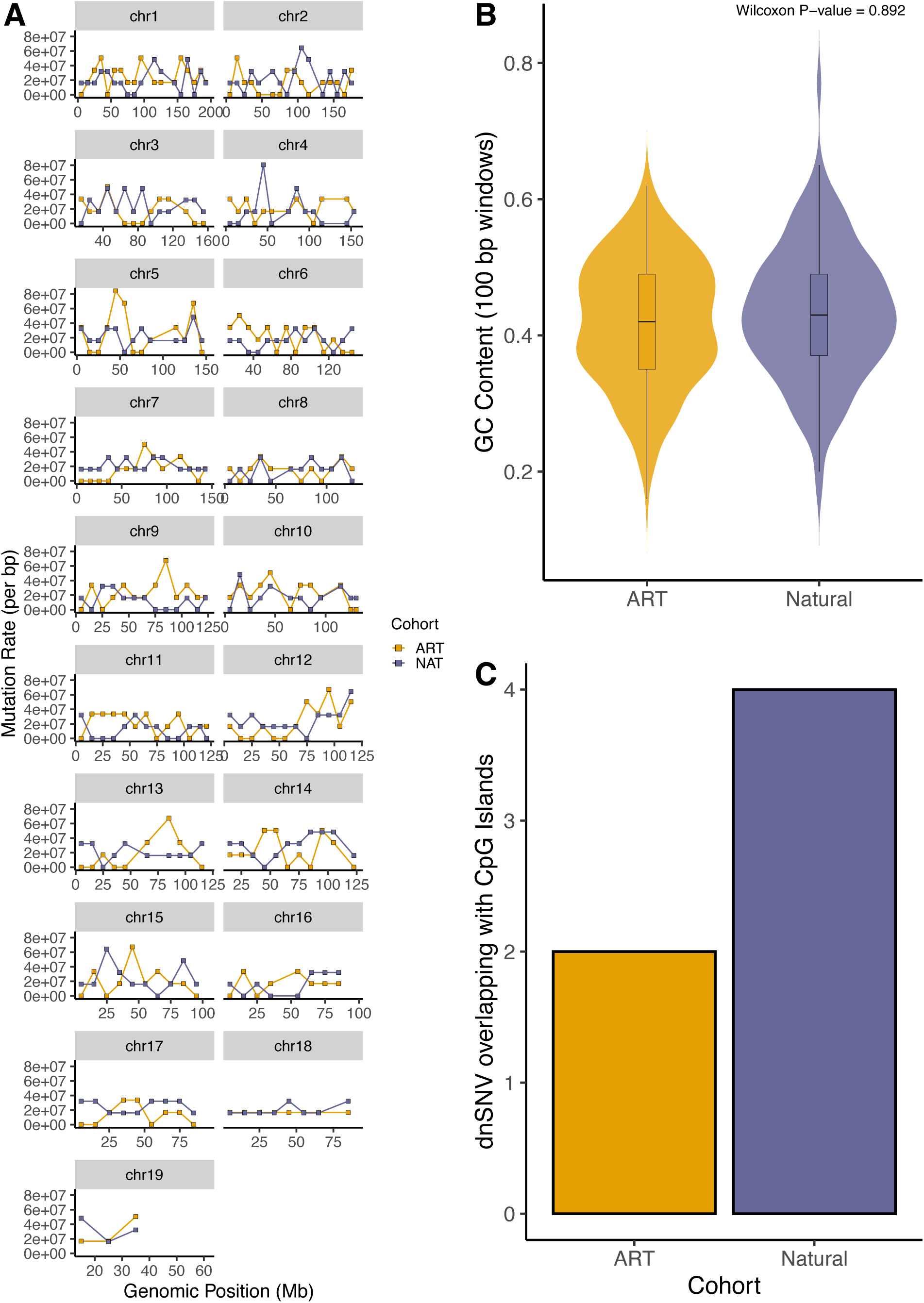
(A)Genomic distribution of dnSNVs in ART-derived (orange) and naturally conceived (purple) cohorts. Mutation counts were aggregated in 10 Mb windows across the autosomal genome. (B) Violin plots depicting the distribution of GC content in 100 bp windows centered on dnSNVs identified in both cohorts. Embedded boxplots represent the interquartile range and median GC content. (C) Bar plot showing the proportion of dnSNVs in ART-and natural-born samples that overlap CpG islands. There is no cohort-level difference (Fisher’s exact test, P = 0.6862).

**Supplementary Figure 7.**
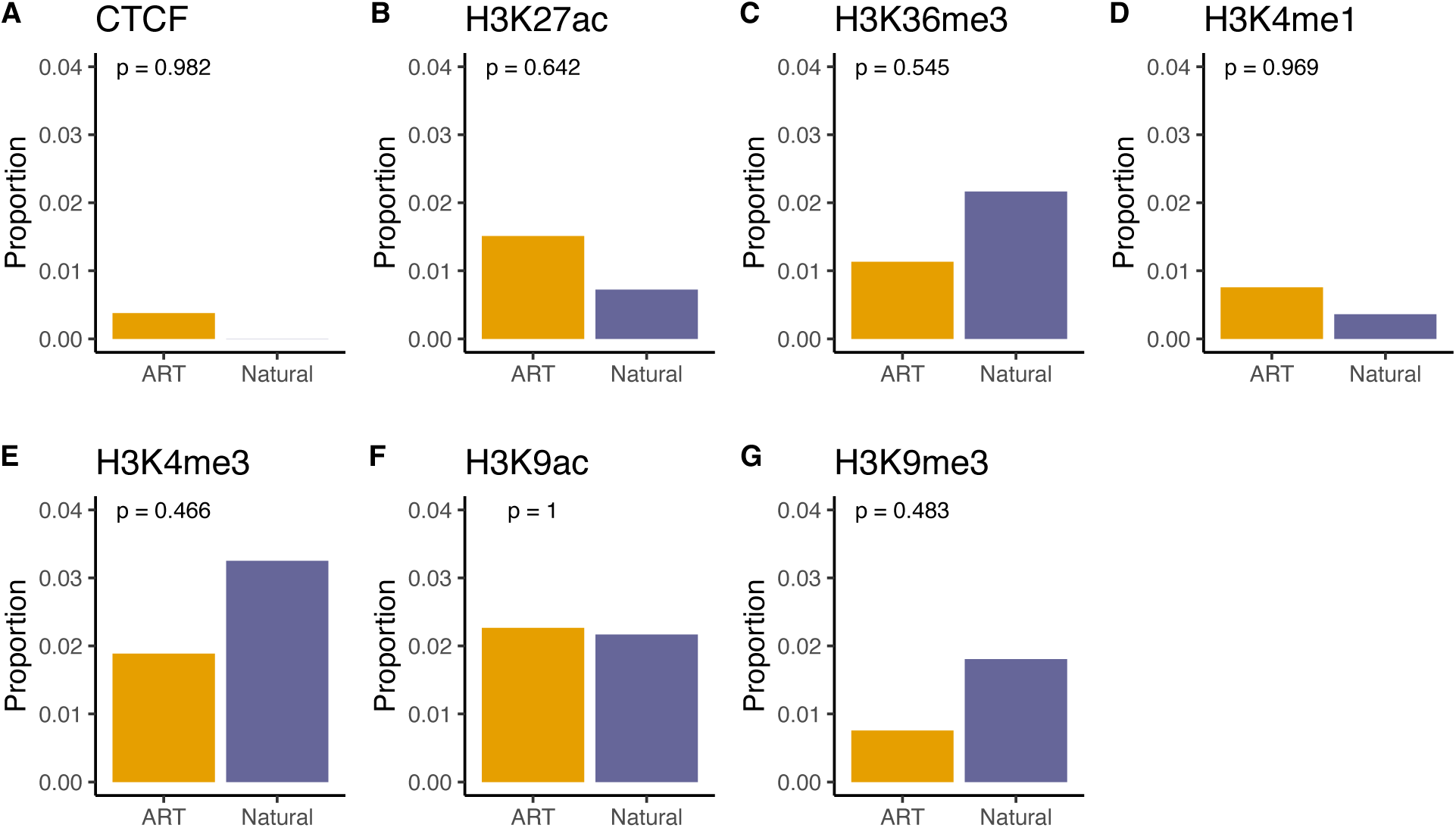
Proportion of dnSNVs in ART- and naturally-born samples that overlap ChIP-seq peaks associated with (A) CTCF binding or (B-G) various histone modifications in mESCs. P-values were computed from Chi-square tests on 1 degree of freedom. ChIP-seq data are from the Bruce4ES dataset released with the mouse ENCODE project.

**Supplementary Figure 8.**
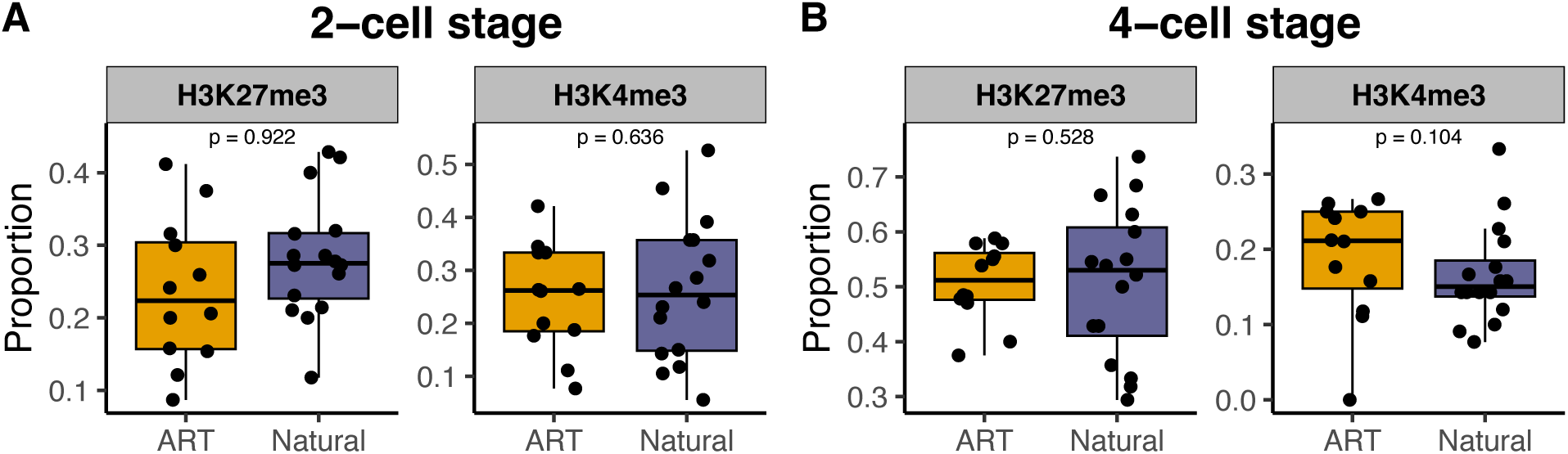
Proportion of dnSNVs in ART- and naturally-born samples that overlap H3K27me3 and H3K4me3 ChIP-seq peaks in 2-cell stage (A) and 4-cell stage (B) mouse embryos. ChIP-seq data are from a published source (54).

**Supplementary Figure 9.**
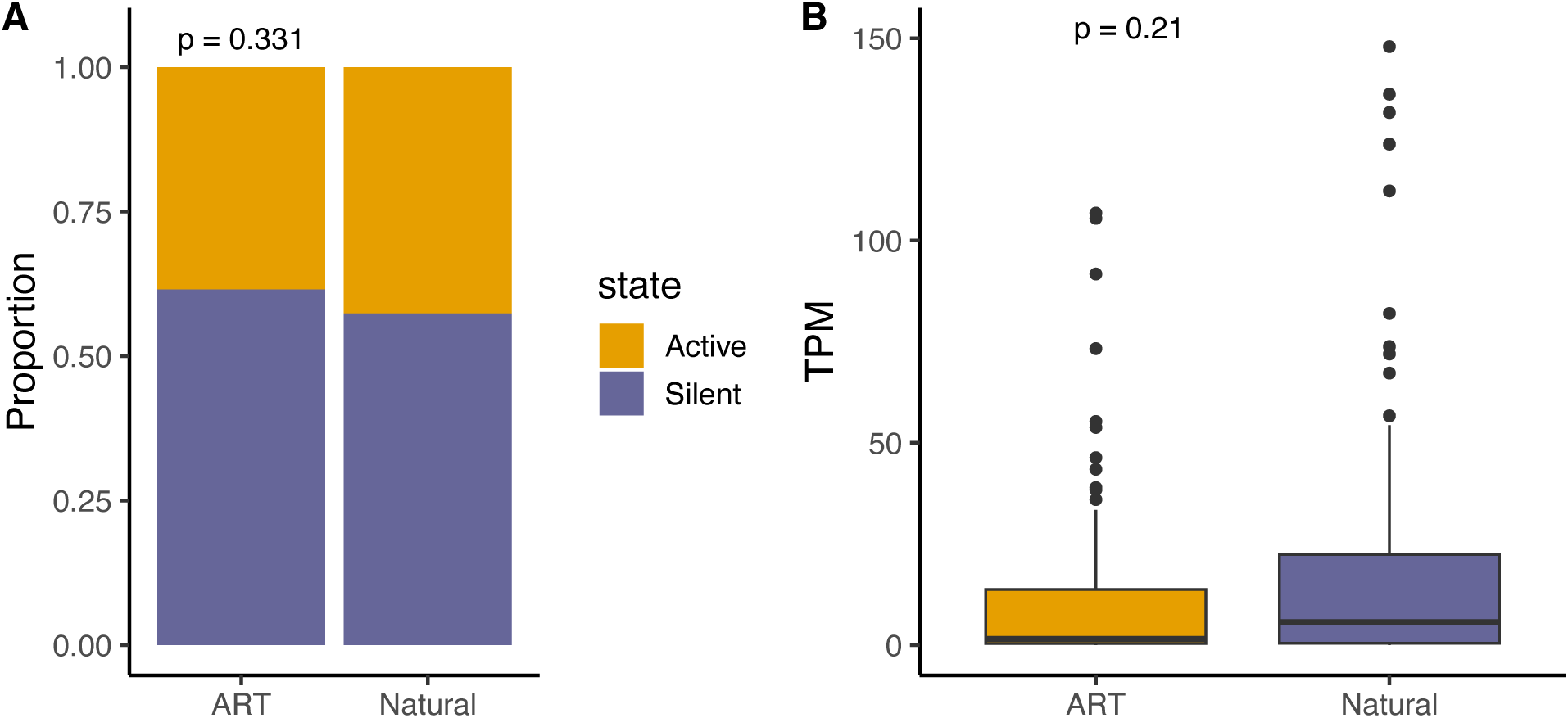
No difference in dnSNV distribution between ART- and natural-born progeny with respect to the transcriptional activity of neighboring genes in C57BL/6J mouse ESCs. dnSNVs were intersected with neighboring genes, with a 2.5kb allowance upstream and downstream of the gene start and end coordinates, respectively. Both (A) the proportion of dnSNVs near genes that are expressed (Active) versus silenced (Silent) and (B) the overall transcript abundance of active genes expressed as transcripts per million (TPM) are indistinguishable between cohorts. Significance was assessed by Chi-square test (A) or Wilcox rank sum test (B).

**Supplementary Figure 10.**
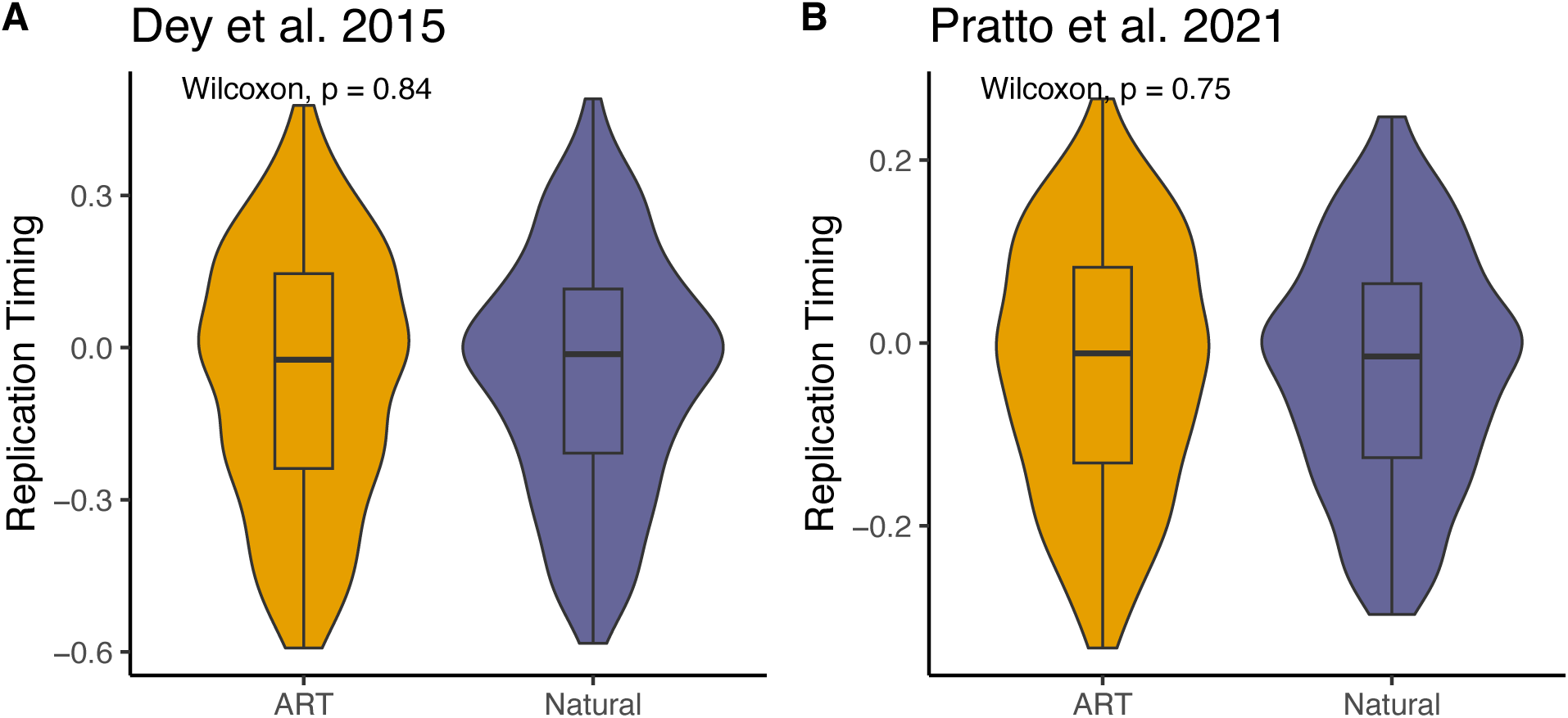
No difference in replication timing of dnSNVs in ART-versus natural-born mice. Replication timing estimates were based on published Repli-seq datasets for mESCs (57, 58). Replication timing estimates are plotted as violin plots with inset boxplots indicating median (thick black line) and interquartile range (box height). Lack of cohort-level differences in replication timing were affirmed using Wilcoxon Rank Sum tests.

**Supplementary Figure 11:**
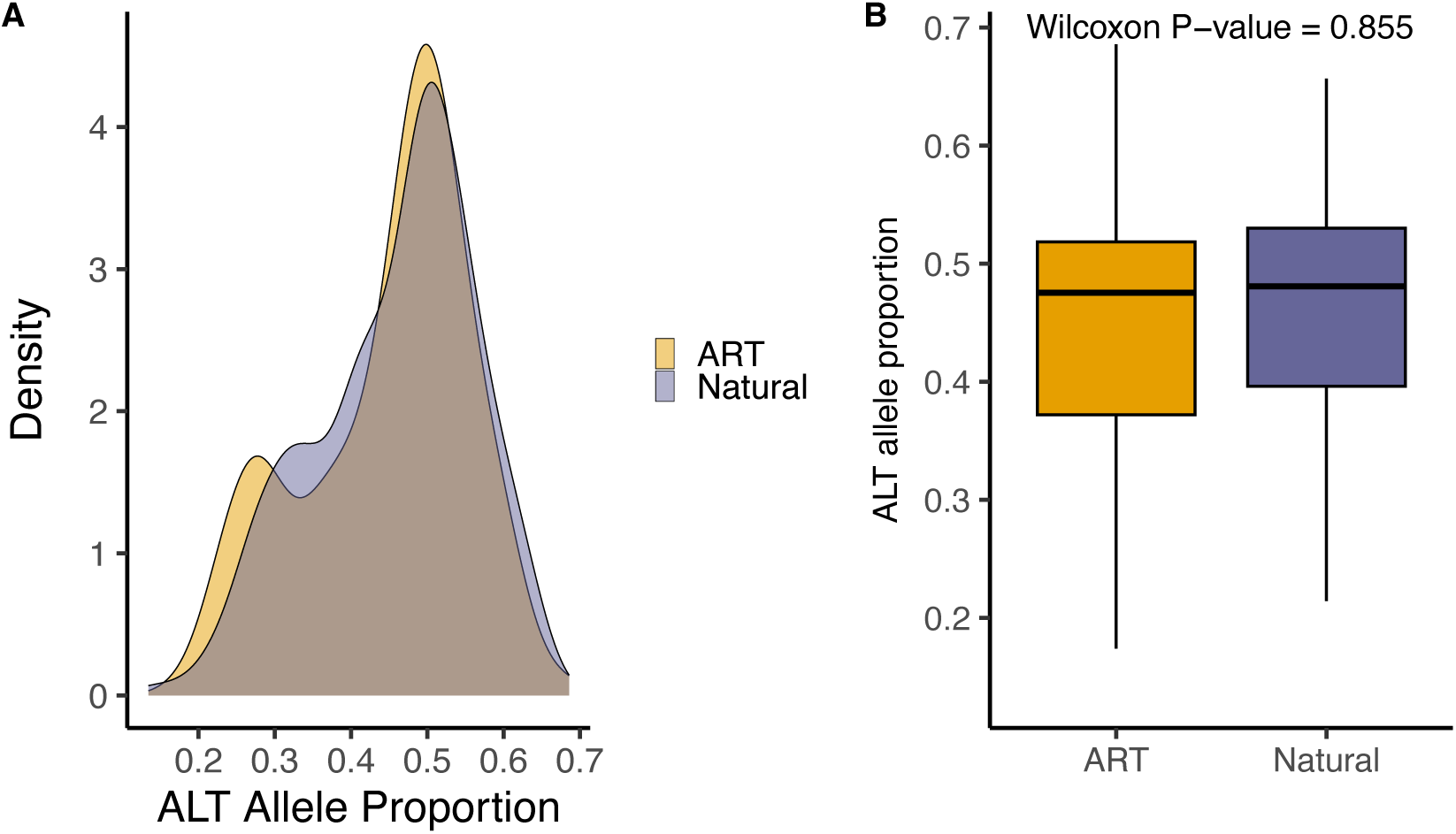
Distribution of ALT allele proportion by cohort. (A) Density plot showing the distribution of ALT allele proportions for dnSNVs in each cohort. (B) Boxplot comparing ALT allele proportions between cohorts.

## SUPPLEMENTARY TABLES

**Supplementary Table 1:** Sequencing metrics for all sequenced samples.

**Supplementary Table 2:** De novo single nucleotide variants in G2 samples using merged data.

**Supplementary Table 3:** De novo single nucleotide variants in G2 samples using Seq1 data.

**Supplementary Table 4:** De novo single nucleotide variants in G2 samples using Seq2 data.

**Supplementary Table 5:** De novo structural variants in G2 samples.

**Supplementary Table 6:** Blacklist gene set comprised of highly copy number variable genes in the mouse genome

## REFERENCES

1. ESHRE ART Fact Sheet. (2025). Available at: https://www.eshre.eu/Press-Room/Resources/Fact-sheets [Accessed 20 June 2025].

2. CDC, State-Specific ART Surveillance. Assisted Reproductive Technology (ART) (2024). Available at: https://www.cdc.gov/art/php/surveillance-state-specific/index.html [Accessed 20 June 2025].

3. J. Smeenk, et al., ART in Europe, 2019: results generated from European registries by ESHRE. Human Reproduction 38, 2321–2338 (2023).

4. Y. Katagiri, et al., Assisted reproductive technology in Japan: A summary report for 2021 by the Ethics Committee of the Japan Society of Obstetrics and Gynecology. Reproductive Medicine and Biology 23, e12552 (2023).

5. CDC, National ART Summary. Assisted Reproductive Technology (ART) (2024). Available at: https://www.cdc.gov/art/php/national-summary/index.html [Accessed 20 June 2025].

6. C. N. Nugent, A. Chandra, Infertility and Impaired Fecundity in Women and Men in the United States, 2015–2019. National Health Statistics Report 202, 1–19 (2024).

7. M. J. Osterman, B. E. Hamilton, J. A. Martin, A. K. Driscoll, C. P. Valenzuela, Births: Final Data for 2023. National Vital Statistics Reports 74 (2025).

8. E. de Waal, et al., The cumulative effect of assisted reproduction procedures on placental development and epigenetic perturbations in a mouse model. Hum Mol Genet 24, 6975– 6985 (2015).

9. G. Koustas, C. Sjoblom, Minute changes to the culture environment of mouse pre-implantation embryos affect the health of the conceptus. Asian Pacific Journal of Reproduction 5, 287–294 (2016).

10. B. Yu, T. H. Smith, S. L. Battle, S. Ferrell, R. D. Hawkins, Superovulation alters global DNA methylation in early mouse embryo development. Epigenetics 14, 780–790 (2019).

11. M. Ooga, et al., Aberrant histone methylation in mouse early preimplantation embryos derived from round spermatid injection. Biochemical and Biophysical Research Communications 680, 119–126 (2023).

12. J. Barberet, et al., Do assisted reproductive technologies and *in vitro* embryo culture influence the epigenetic control of imprinted genes and transposable elements in children? Human Reproduction 36, 479–492 (2021).

13. C. Choux, et al., The epigenetic control of transposable elements and imprinted genes in newborns is affected by the mode of conception: ART versus spontaneous conception without underlying infertility. Human Reproduction 33, 331–340 (2018).

14. J. Ghosh, C. Coutifaris, C. Sapienza, M. Mainigi, Global DNA methylation levels are altered by modifiable clinical manipulations in assisted reproductive technologies. Clinical Epigenetics 9, 14 (2017).

15. S. Mani, J. Ghosh, C. Coutifaris, C. Sapienza, M. Mainigi, Epigenetic changes and assisted reproductive technologies. Epigenetics 15, 12 (2019).

16. S. E. Håberg, et al., DNA methylation in newborns conceived by assisted reproductive technology. Nat Commun 13, 1896 (2022).

17. S. Song, et al., DNA methylation differences between in vitro- and in vivo-conceived children are associated with ART procedures rather than infertility. Clin Epigenetics 7, 41 (2015).

18. R. M. Rivera, et al., Manipulations of mouse embryos prior to implantation result in aberrant expression of imprinted genes on day 9.5 of development. Hum Mol Genet 17, 1– 14 (2008).

19. G. Giritharan, et al., Effect of in vitro fertilization on gene expression and development of mouse preimplantation embryos. Reproduction 134, 63–72 (2007).

20. S. Hayashi, Mouse Preimplantation Embryos Developed from Oocytes Injected with Round Spermatids or Spermatozoa Have Similar but Distinct Patterns of Early Messenger RNA Expression. Biology of Reproduction 69, 1170–1176 (2003).

21. T. Kohda, Effects of embryonic manipulation and epigenetics. J Hum Genet 58, 416–420 (2013).

22. K. Tan, et al., Dynamic integrated analysis of DNA methylation and gene expression profiles in in vivo and in vitro fertilized mouse post-implantation extraembryonic and placental tissues. Mol Hum Reprod 22, 485–498 (2016).

23. V. K. Cortessis, et al., Comprehensive meta-analysis reveals association between multiple imprinting disorders and conception by assisted reproductive technology. Journal of Assisted Reproduction and Genetics 35, 943 (2018).

24. H. Hattori, et al., Association of four imprinting disorders and ART. Clinical Epigenetics 11, 21 (2019).

25. S. D. McDonald, et al., Preterm birth and low birth weight among in vitro fertilization singletons: a systematic review and meta-analyses. Eur J Obstet Gynecol Reprod Biol 146, 138–148 (2009).

26. S. Palomba, R. Homburg, S. Santagni, G. B. La Sala, R. Orvieto, Risk of adverse pregnancy and perinatal outcomes after high technology infertility treatment: a comprehensive systematic review. Reprod Biol Endocrinol 14, 76 (2016).

27. B. Luke, et al., The risk of birth defects with conception by ART. Hum Reprod 36, 116–129 (2021).

28. R. J. Hart, L. A. Wijs, The longer-term effects of IVF on offspring from childhood to adolescence. Frontiers in Reproductive Health 4, 1045762 (2022).

29. C. J. Bean, Fertilization in vitro increases non-disjunction during early cleavage divisions in a mouse model system. Human Reproduction 17, 2362–2367 (2002).

30. M. Bonduelle, Prenatal testing in ICSI pregnancies: incidence of chromosomal anomalies in 1586 karyotypes and relation to sperm parameters. Human Reproduction 17, 2600– 2614 (2002).

31. Y.-L. Lee, et al., The rate of de novo structural variation is increased in in vitro–produced offspring and preferentially affects the paternal genome. Genome Res. 33, 1455–1464 (2023).

32. M. Zamani Esteki, et al., In vitro fertilization does not increase the incidence of de novo copy number alterations in fetal and placental lineages. Nat Med 25, 1699–1705 (2019).

33. L. Caperton, et al., Assisted reproductive technologies do not alter mutation frequency or spectrum. Proc. Natl. Acad. Sci. U.S.A. 104, 5085–5090 (2007).

34. C. Wang, et al., Association of assisted reproductive technology, germline de novo mutations and congenital heart defects in a prospective birth cohort study. Cell Res 31, 919–928 (2021).

35. K. S. Ruth, et al., Genetic insights into biological mechanisms governing human ovarian ageing. Nature 596, 393–397 (2021).

36. K. D. Makova, R. C. Hardison, The effects of chromatin organization on variation in mutation rates in the genome. Nat Rev Genet 16, 213–223 (2015).

37. C. Chen, H. Qi, Y. Shen, J. Pickrell, M. Przeworski, Contrasting Determinants of Mutation Rates in Germline and Soma. Genetics 207, 255–267 (2017).

38. E. Vanneste, et al., Chromosome instability is common in human cleavage-stage embryos. Nat Med 15, 577–583 (2009).

39. C. E. Currie, et al., The first mitotic division of human embryos is highly error prone. Nat Commun 13, 6755 (2022).

40. O. Konstantogianni, et al., Culture of Human Embryos at High and Low Oxygen Levels. J Clin Med 13, 2222 (2024).

41. J. J. Gille, C. G. van Berkel, H. Joenje, Mutagenicity of metabolic oxygen radicals in mammalian cell cultures. Carcinogenesis 15, 2695–2699 (1994).

42. R. G. Bristow, R. P. Hill, Hypoxia, DNA repair and genetic instability. Nat Rev Cancer 8, 180–192 (2008).

43. C. Zhao, et al., Single-cell multi-omics of human preimplantation embryos shows susceptibility to glucocorticoids. Genome Res. 32, 1627–1641 (2022).

44. A. Garretson, L. Blanco-Berdugo, A. Roberts, B. L. Dumont, Benchmarking Genomic Variant Calling Tools in Inbred Mouse Strains: Recommendations and Considerations. [Preprint] (2025). Available at: https://www.biorxiv.org/content/10.1101/2025.05.28.656711v1 [Accessed 20 June 2025].

45. A. Uchimura, et al., Germline mutation rates and the long-term phenotypic effects of mutation accumulation in wild-type laboratory mice and mutator mice. Genome Res 25, 1125–1134 (2015).

46. S. J. Lindsay, R. Rahbari, J. Kaplanis, T. Keane, M. E. Hurles, Similarities and differences in patterns of germline mutation between mice and humans. Nat Commun 10, 4053 (2019).

47. L. A. Bergeron, et al., Evolution of the germline mutation rate across vertebrates. Nature 615, 285–291 (2023).

48. E. López-Cortegano, et al., The rate and spectrum of new mutations in mice inferred by long-read sequencing. Genome Res. 35, 43–54 (2025).

49. E. D. Pleasance, et al., A comprehensive catalogue of somatic mutations from a human cancer genome. Nature 463, 191–196 (2010).

50. L. B. Alexandrov, et al., The repertoire of mutational signatures in human cancer. Nature 578, 94–101 (2020).

51. E. López-Cortegano, et al., Variation in the Spectrum of New Mutations among Inbred Strains of Mice. Mol Biol Evol 41, msae163 (2024).

52. W. Reik, W. Dean, J. Walter, Epigenetic reprogramming in mammalian development. Science 293, 1089–1093 (2001).

53. F. Yue, et al., A comparative encyclopedia of DNA elements in the mouse genome. Nature 515, 355–364 (2014).

54. X. Liu, et al., Distinct features of H3K4me3 and H3K27me3 chromatin domains in pre-implantation embryos. Nature 537, 558–562 (2016).

55. L. Moore, et al., The mutational landscape of human somatic and germline cells. Nature 597, 381–386 (2021).

56. J. A. Stamatoyannopoulos, et al., Human mutation rate associated with DNA replication timing. Nat Genet 41, 393–395 (2009).

57. F. Pratto, et al., Meiotic recombination mirrors patterns of germline replication in mice and humans. Cell 184, 4251–4267 (2021).

58. S. S. Dey, L. Kester, B. Spanjaard, M. Bienko, A. van Oudenaarden, Integrated genome and transcriptome sequencing of the same cell. Nat Biotechnol 33, 285–289 (2015).

59. J. R. Belyeu, et al., De novo structural mutation rates and gamete-of-origin biases revealed through genome sequencing of 2,396 families. Am J Hum Genet 108, 597–607 (2021).

60. O. I. Garcia-Salinas, et al., The impact of ancestral, genetic, and environmental influences on germline de novo mutation rates and spectra. Nat Commun 16, 4527 (2025).

61. T. Hassold, P. Hunt, To err (meiotically) is human: the genesis of human aneuploidy. Nat Rev Genet 2, 280–291 (2001).

62. M. Hansen, J. J. Kurinczuk, E. Milne, N. de Klerk, C. Bower, Assisted reproductive technology and birth defects: a systematic review and meta-analysis. Hum Reprod Update 19, 330–353 (2013).

63. M. J. Davies, et al., Reproductive Technologies and the Risk of Birth Defects. New England Journal of Medicine 366, 1803–1813 (2012).

64. Y. Yamauchi, et al., Loss of mouse Y chromosome gene Zfy1 and Zfy2 leads to spermatogenesis impairment, sperm defects, and infertility. Biol Reprod 106, 1312–1326 (2022).

65. R. A. Taft, M. Davisson, M. V. Wiles, Know thy mouse. Trends Genet 22, 649–653 (2006).

66. N. Dukler, M. R. Mughal, R. Ramani, Y.-F. Huang, A. Siepel, Extreme purifying selection against point mutations in the human genome. Nat Commun 13, 4312 (2022).

67. S. Chen, Ultrafast one-pass FASTQ data preprocessing, quality control, and deduplication using fastp. iMeta 2, e107 (2023).

68. T. Yun, et al., Accurate, scalable cohort variant calls using DeepVariant and GLnexus. Bioinformatics 36, 5582–5589 (2021).

69. W. J. Kent, et al., The Human Genome Browser at UCSC. Genome Res. 12, 996–1006 (2002).

70. P. Danecek, et al., The variant call format and VCFtools. Bioinformatics 27, 2156–2158 (2011).

71. M. Díaz-Gay, et al., Assigning mutational signatures to individual samples and individual somatic mutations with SigProfilerAssignment. Bioinformatics 39, btad756 (2023).

72. P. Cingolani, et al., A program for annotating and predicting the effects of single nucleotide polymorphisms, SnpEff. Fly (Austin) 6, 80–92 (2012).

73. J. A. Stamatoyannopoulos, et al., An encyclopedia of mouse DNA elements (Mouse ENCODE). Genome Biology 13, 418 (2012).

74. T. Rausch, et al., DELLY: structural variant discovery by integrated paired-end and split-read analysis. Bioinformatics 28, i333–i339 (2012).

75. X. Chen, et al., Manta: rapid detection of structural variants and indels for germline and cancer sequencing applications. Bioinformatics 32, 1220–1222 (2016).

76. A. C. English, V. K. Menon, R. A. Gibbs, G. A. Metcalf, F. J. Sedlazeck, Truvari: refined structural variant comparison preserves allelic diversity. Genome Biology 23, 271 (2022).

77. J. R. Belyeu, et al., Samplot: a platform for structural variant visual validation and automated filtering. Genome Biology 22, 161 (2021).

78. W. McLaren, et al., The Ensembl Variant Effect Predictor. Genome Biology 17, 122 (2016).

